# THRESHOLD: A Comprehensive Transcriptomic Analysis Tool for Evaluating Gene Saturation and Impact in Disease Progression

**DOI:** 10.1101/2024.10.20.619322

**Authors:** Finán Gammell, Jennifer Li, Christopher Elco, Jessica Plavicki, Alper Uzun

## Abstract

Gene expression studies serve as a foundational tool in molecular biology, providing insights into developmental, physiological, and pathological processes. Variations in gene expression can indicate disease states, which are vital in understanding disease progression, subtype manifestations, and identifying therapeutic targets based on detailed expression patterns. To effectively investigate gene expression patterns, especially in large datasets, a robust and precise analysis tool is crucial. In response to this critical analytical need, we developed THRESHOLD, a novel tool that goes beyond traditional gene expression analysis by introducing the concept of gene saturation. Unlike conventional methods that focus on absolute expression levels or binary differential expression, THRESHOLD quantifies the consistency of gene expression across patients, revealing co-regulation patterns that may otherwise be overlooked. This novel metric offers a unique perspective on gene expression patterns by highlighting consistent trends across patient samples, which are critical for understanding disease mechanisms and stratifying patients based on molecular signatures. The tool offers several features, including user-defined parameters, statistical comparisons, and interactive data visualization. THRESHOLD has uncovered compelling insights into disease progression using TCGA Cancer Datasets. For instance, bladder urothelial carcinoma demonstrated increasing upregulated gene saturation in progressive cancer stages (p < 0.00001). Moreover, THRESHOLD identified heightened gene saturation in patients with earlier onset of prostate adenocarcinoma (p < 0.0001) and revealed a critical fusion transcript, SLC45A2-AMACR, implicated in prostate adenocarcinoma progression, recurrence, and metastasis. Additionally, novel biomarkers and potential candidates for drug therapies were identified through protein-protein interaction networks and functional analyses of saturation data in colon adenocarcinoma and breast invasive carcinoma. Collectively, THRESHOLD advances our understanding of patient stratification and molecular signatures by offering a more detailed view of gene expression dynamics. The THRESHOLD tool is publicly available at: https://github.com/alperuzun/THRESHOLD.

## Introduction

Researchers and clinicians’ ability to accurately diagnose and treat diseases is aided by their capacity to understand disease heterogeneity (1) as it affects to disease etiology and progression. Cancer is a complex disease and clinically challenging to treat due to the numerous pathways through which carcinogenesis can occur and the molecular variation within cancer subtypes (2). Therefore, it is highly valuable for researchers and clinicians to identify significant differences in gene expression. These differences can help predict disease progression and determine whether a tumor is likely to respond to or resist various therapeutic approaches.

Cancer remains a leading cause of global mortality with incidence rates increasing at an alarming pace. New cancer cases in the United States have increased 30.8% from around 1,500,000 in 2010 to 2,000,000 in 2024 (3, 4). Changes in new cancer cases cannot be explained by population changes (5, 6). The molecular complexity of cancer coupled with the increase in cancer incidence creates a critical need for new analytical approaches for analyzing clinical samples in a manner that aids in the development of new and affective therapeutic approaches (7). Examining the subtle biological pathways through which illnesses arise and manifest can help clarify these complexities.

The field of genomics offers promise in healthcare in its capacity to aid in early detection of diseases, patient stratification, and personalized medicine, with gene expression analysis driving this innovation. Stratified analyses of upregulated and downregulated genes can shed light on potential pathways and therapeutic targets across patient groups of interest. The patterns in which genes are expressed can unveil significant insights into disease mechanisms, guide therapeutic responses, and help interpret complex biological interactions. Currently, differential gene expression (DGE) analysis is utilized as an existing computational approach to evaluate gene expression and compare differences in gene expression between across relevant groups or conditions (8).

Although THRESHOLD offers a novel approach by focusing on gene saturation and consistency in gene expression across patient cohorts, several existing tools can be considered the closest in terms of application, though their methodologies differ. DESeq2 (9) is a widely used differential gene expression (DGE) analysis tool that emphasizes identifying statistically significant changes in gene expression across conditions, but it may overlook consistent gene expression patterns across individuals— a key aspect that THRESHOLD is designed to capture. Similarly, edgeR (10) is another DGE tool that excels in detecting differentially expressed genes in RNA-seq datasets but, like DESeq2, focuses primarily on the magnitude of expression changes rather than consistent gene patterns. Limma (11) is a versatile tool that uses linear models for differential expression analysis of microarray and RNA-seq data, providing valuable insights into gene expression profiling but lacking the focus on gene expression consistency that THRESHOLD offers. Additionally, while CIBERSORT (12) is not a DGE tool, it focuses on deconvoluting immune cell populations from bulk RNA-seq data, and its statistical framework for analyzing gene expression across patient samples offers a useful point of comparison with THRESHOLD’s capabilities in evaluating gene expression patterns across populations. Despite these differences, these tools represent the most relevant comparisons in terms of analyzing gene expression in large datasets.

DGE’s underlying calculations rely on the identification of genes that demonstrate statistically significant changes in expression across patients as compared to some standard, revealing which genes are upregulated or downregulated in response to specific conditions. This approach provides value for detecting major expression shifts and associating them with biological processes or relevant disease states. However, DGE analysis is inherently limited in its ability to capture the broader, more distinct gene expression patterns by patients that may be crucial for understanding complex disease pathology. Specifically, DGE analysis emphasizes the magnitude of expression changes and relies on grouped averages which can be skewed by individual patients, potentially overlooking genes that play consistent roles across individual patients but do not exhibit as significant changes in expression. Analyses that stress the consistency of gene expression ranks across a patient cohort can aid in bridging this gap, helping to identify potentially overlooked disease mechanisms and facilitate representative patient stratification.

To address DGE’s limitations, and complement its analyses through a distinct form of calculation, we have developed a new tool called THRESHOLD, which offers a novel computational analysis to visualize both overarching and detailed patterns of gene expression across large patient samples.

THRESHOLD is a tool for analysis of transcriptomic data across large samples of patients to understand the cohesion of the most upregulated/downregulated genes in a given disease. The tool generates a novel metric called saturation, which quantifies the proportion of genes occurring frequently at similar ranks of relative upregulation. THRESHOLD offers several user-inputted parameters that allow researchers to shape their analyses to the desired saturation type, restriction factors and rank type. Additionally, THRESHOLD offers in-tool statistical analyses to help understand significant stratification between patterns of gene expression. The tool outputs an interactive graph of saturation permitting the calculation of specific saturation thresholds and most saturated genes.

THRESHOLD includes two distinct analytical features, incremental saturation, and overall saturation. Incremental saturation is designed to articulate the step-by-step changes in gene expression patterns by nth rank, whereas overall saturation offers a comprehensive view of changes in gene expression up to a specific nth rank. In addition, THRESHOLD can generate gene ranks based on high to low (most upregulated) or low to high (most downregulated) expression. This calibration allows researchers to easily investigate both the most upregulated or downregulated genes depending on their needs, expanding THRESHOLD’s utility. The primary aim of developing THRESHOLD is to facilitate our understanding of the multifaceted transcriptomic landscape. To that end, it plays an essential role in several key domains: **(1) Disease Sub-stratification:** Beyond merely classifying diseases based on observable symptoms or common biomarkers, THRESHOLD provides a detailed molecular perspective, identifying unique genetic markers within different patient groups and enabling a more refined and precise disease categorization. **(2) Drug target Identification:** THRESHOLD can be used to identify genes that are highly upregulated or downregulated across diverse patient populations. The highly upregulated or downregulated genes occurring across large numbers of patients are likely indicative of genes critical to the disease’s pathology and carcinogenesis. Thus, THRESHOLD’s analyses can aid in the identification of new potential therapeutic targets. **(3) Biomarker Discovery:** In conditions such as cancer, where prompt diagnosis is crucial, THRESHOLD offers valuable assistance. THRESHOLD helps researchers identify genes that are frequently overexpressed, suggesting their possible role as indicators of disease and aiding in the development of early detection methods. Together, the THRESHOLD serves as a useful addition to genomics research by providing insights into disease mechanisms and supporting the advancements in personalized medicine.

## Materials and Methods

### Data Requirements

THRESHOLD requires the input of a file of patient transcriptomic data with gene expression data in the form of z-scores, comparing expression against a control population, or percentiles, ranking expression relatively within an individual patient’s transcriptomic profile.

The input file for THRESHOLD should be in a tab-delimited (‘\t’) format in a text (.txt) file. Each row should represent a patient sample, and each column should correspond to a specific gene or biomarker. The first row of the file should contain the header, including a required “Hugo_Symbol” (HUGO Gene Nomenclature Committee) title for the first column, a blank space on the second column, and the relevant patient identifier for each successive column. The first column (“Hugo_Symbol”) should include unique identifiers of each gene, the second column should remain empty, while each successive column should include expression data by each relevant patient. Missing expression data should be represented by “NA.”

### Saturation Analysis Methods

THRESHOLD offers two primary functionalities: “Overall Saturation” and “Incremental Saturation.” The former offers a holistic perspective on gene expression by considering the saturation of all genes up to and including the nth level, while the latter focuses on saturation at each specific nth gene level. Both are original metrics developed to facilitate THRESHOLD’s novel outputs.

#### Overall Saturation

The saturation of all the genes up to and including the nth level. Quantifies the count of genes up to the nth level that exceed the inputted restriction level, divided by the total count of genes up to and including the nth level, Figure 2.

Quantitatively, this is represented by the sum of each unique element in the multiset A’s non-negative greater value between the element count in multiset A minus the restriction level divided by the number of elements in the multiset A. Formula for overall saturation was presented in Figure 1.

**Figure 1.**
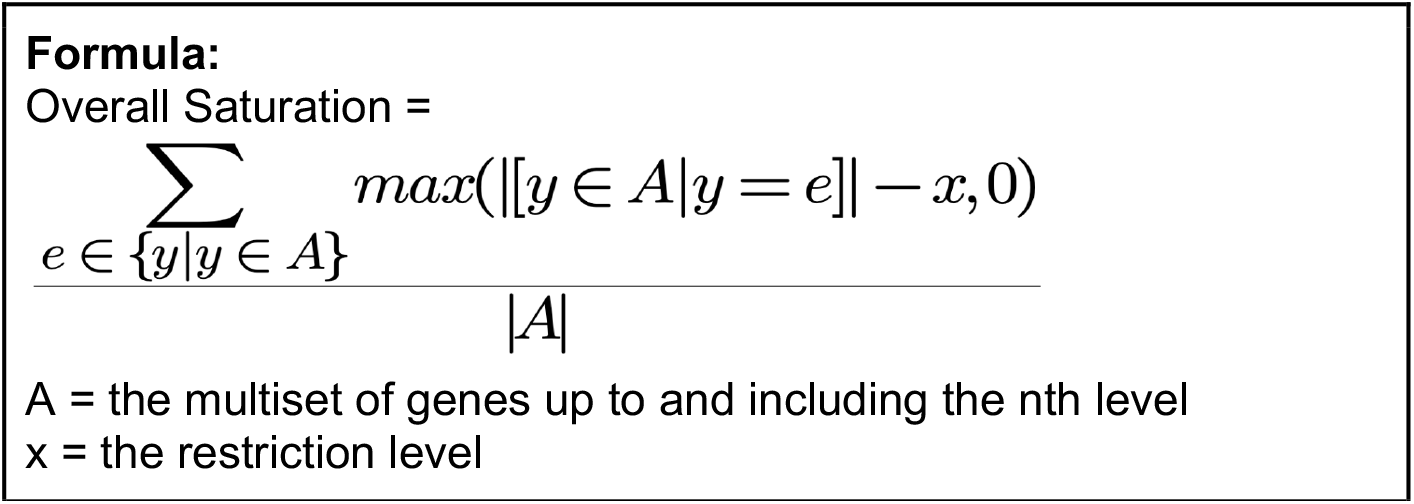
Formula for overall saturation.

### Visual Example

***Find the overall saturation up to the 3rd level (n = 3) given a restriction level of 1***:

#### Incremental Saturation

The saturation only at the incremental nth gene level. Quantifies the count of genes at the nth level that exceed the inputted restriction level up to and including that level, divided by the total count of genes at the nth level, Figure 4.

Quantitatively, this is represented by the sum of each unique element in the multiset B’s non-negative lesser value between the difference in the element count in multiset A minus the restriction level and the element count in multiset B divided by the number of elements in multiset B. Formula for overall saturation was presented in Figure 3.

**Figure 2.**
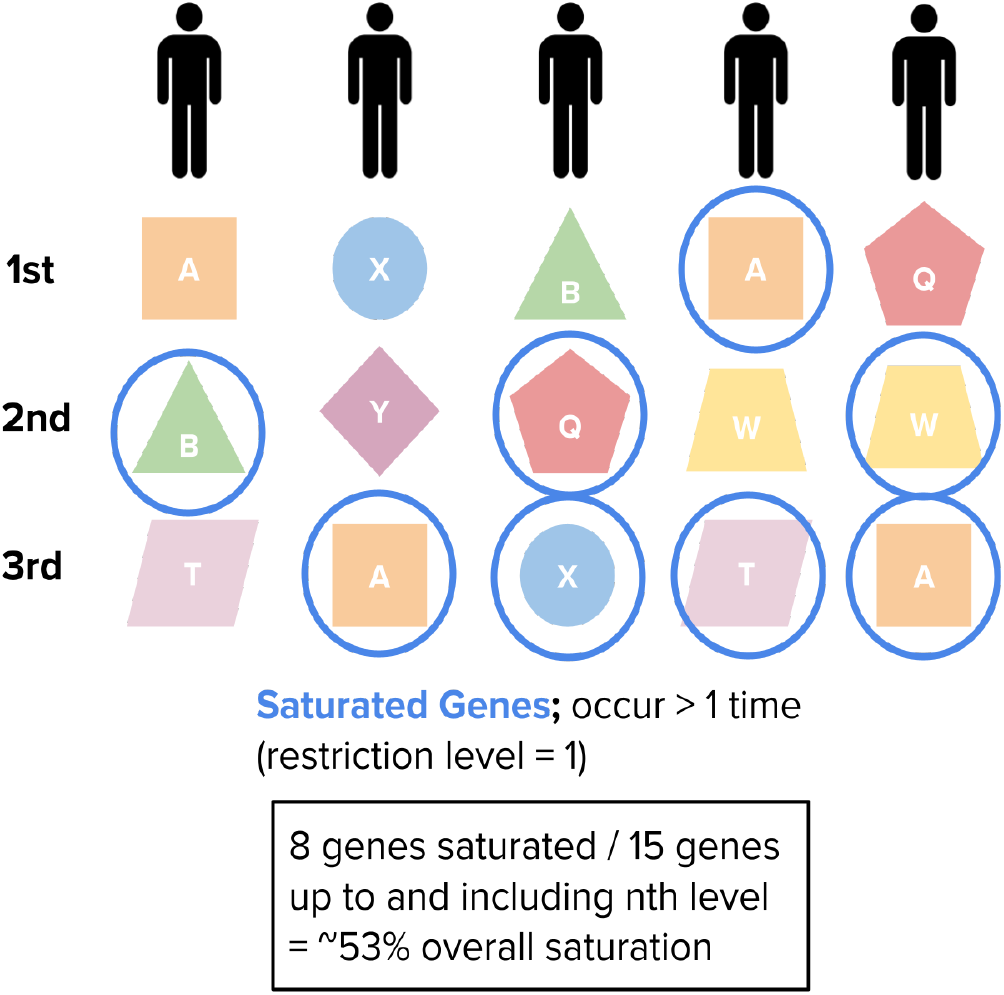
Overall Saturation Visual Example. Genes are ranked by most relatively upregulated for each patient group, represented by each column of shapes. Given a restriction level of one, any gene that occurs beyond the first occurrence by nth level is considered saturated. For overall saturation, saturation is calculated by the proportion of saturated genes at all levels up to and including a given nth level, n = 3 in this case.

**Figure 3.**
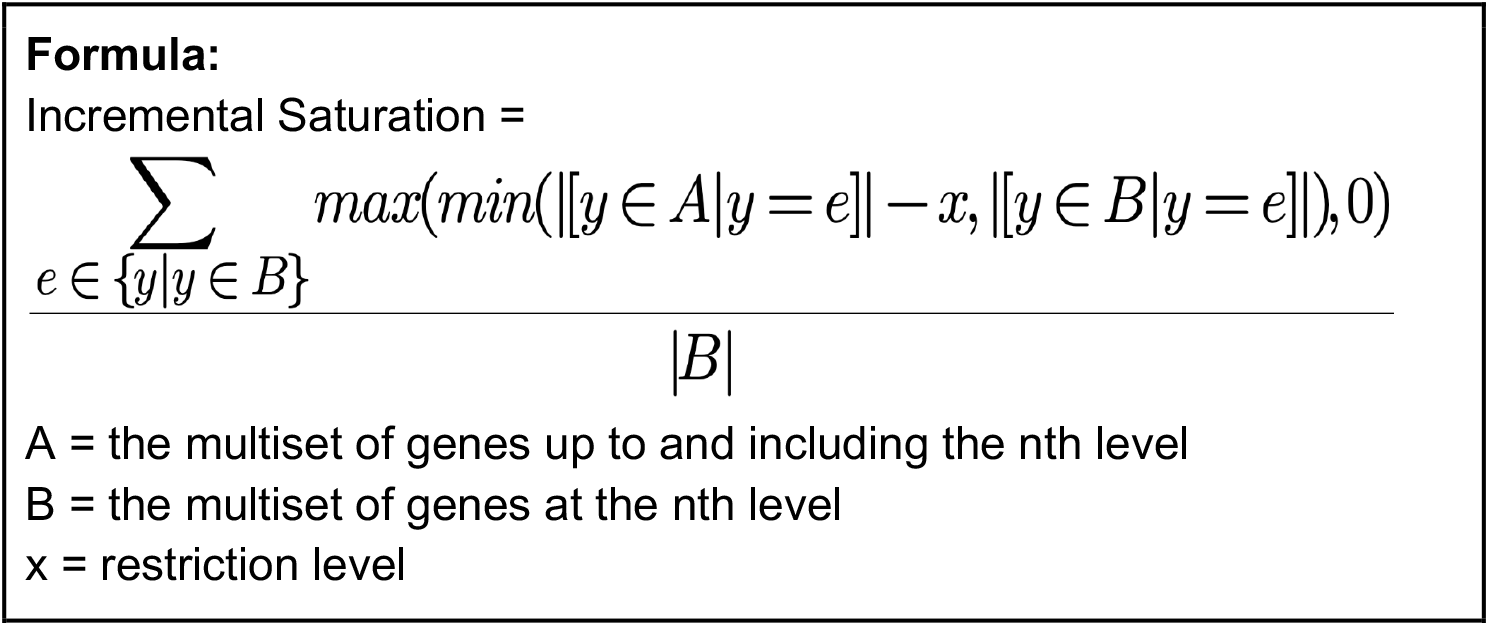
Formula for incremental saturation.

**Figure 4.**
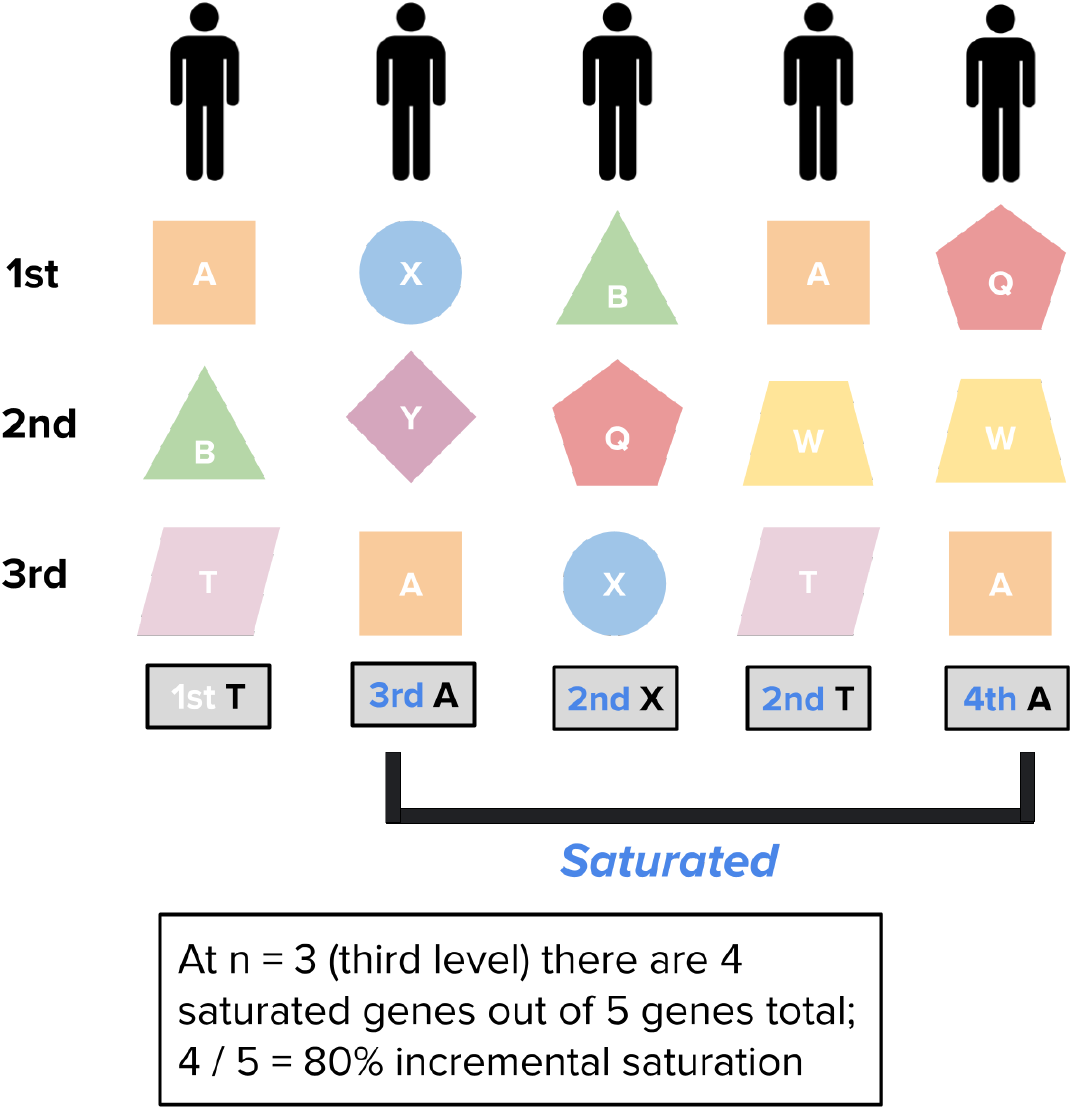
Incremental Saturation Visual Example. Genes are ranked by most relatively upregulated for each patient group, represented by each column of shapes. Given a restriction level of one, any gene that occurs beyond the first occurrence by nth level is considered saturated. For incremental saturation, saturation is calculated by the proportion of saturated genes specifically present at the given nth level, n = 3 in this case.

### Visual Example

***Find the incremental saturation of the 3rd level (n = 3) given a restriction level of 1***:

### Software and Algorithm

The THRESHOLD software was built using Python and Java and leveraged dependencies including NumPy, matplotlib, SciPy, and PyQt6 to support calculations, visualizations, statistics, and GUI display respectively. The core algorithm relies on a novel statistical metric developed called “Saturation”, which systematically analyzes gene expression data to detect highly expressed genes shared across different patient datasets.

To demonstrate the application of THRESHOLD, gene expression data from TCGA (13) datasets accessed via UCSC Xena (14) were used, and clinical features from cBioPortal (15) were incorporated for further comparative analyses. The datasets spanned several cancers including bladder urothelial carcinoma, prostate adenocarcinoma, colon adenocarcinoma, and breast invasive carcinoma, incorporating gene expression profiles of patients. THRESHOLD requires the input of a file of patient sample transcriptomic data with gene expression data in the form of z-scores or percentiles comparing expression against a control population or ranked relatively within an individual patient’s expression. Such data is publicly available in many TCGA datasets via UCSC Xena, cBioPortal, and more, Figure 5.

**Figure 5.**
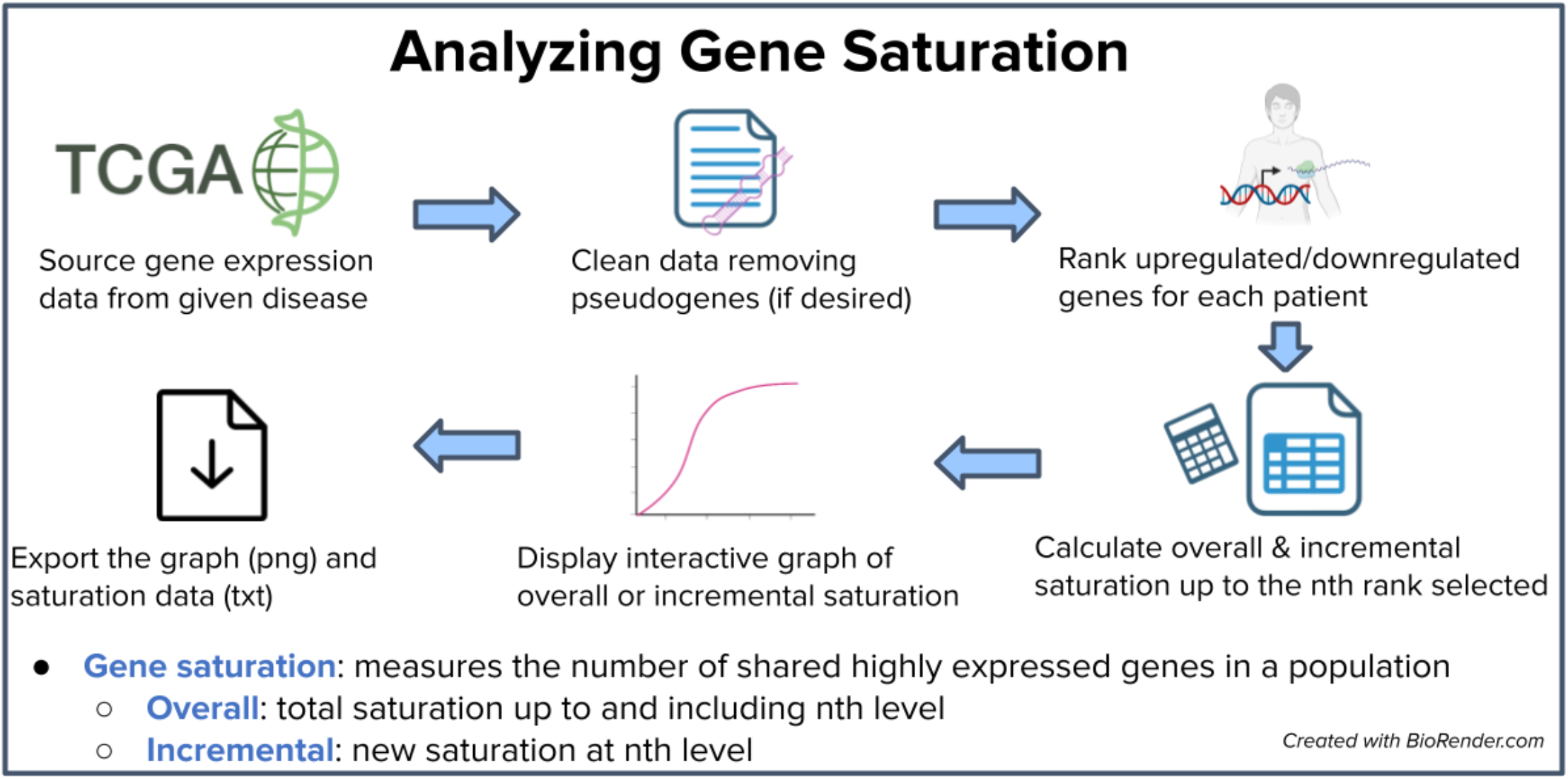
Overview of GUI Workflow. An outline of THRESHOLD’s workflow, specifically how data is organized in order to calculate and understand gene saturation. *Created with BioRender*.*com*^*11*^

### Data Preprocessing

Once uploaded to the THRESHOLD GUI, the data will automatically be cleaned to ensure the most relevant results. This will include removing pseudogenes including non-coding RNA such as snoRNA, sniRNA, snRNA, and microRNA. The removed pseudogenes will be exported to a resulting .txt file to validate appropriate removal. If a user wants to keep these pseudogenes, they can also select to include them. Gene expressions is then ranked for each patient by most upregulated or downregulated depending on user-inputted selection. These ranks are recorded as JSON data for each patient. This rank leverages the z-score/percentile value assigned to each gene from the given dataset. Once saturation values are calculated for each level up to the user-inputted request, the saturation data is saved as a downloadable .txt file and displayed within an interactive graph. Users can calculate specific saturation values at given levels, and also find when a saturation threshold is met. Both the graph and the aforementioned .txt file of the overall saturation data is easily exportable for further analysis or records.

## Comparative Analysis

### Standardization

To compare saturation across datasets, restriction levels can be standardized. Increasing the restriction level by a factor of the data set size increases and produces the same saturation results and can be used for standardization. For example, a data set of 100 patients has a restriction level of 2 genes. To standardize a data set of 200 patients (twice the size), you would need to double the restriction level 2×2 = 4.

It is important to mention that restriction levels in terms of a percentage are universal and do not need further standardization. However, due to round by patient sample size, there may be potentially ±1 gene differences in the restriction level.

### Statistical Analysis

THRESHOLD also facilitates the statistical comparison between two saturation data sets to assess whether there are statistically significant differences between samples to provide additional insights in user analyses. Ensure the files are in the proper, standard format (.txt) as exported by the THRESHOLD tool. The files should have three columns, “Nth Gene Included,” “Incremental Saturation,” and “Overall Saturation.” The file should be the same size, i.e., the same number of rows. The test of choice is a Two-Tailed Unpaired T-Test, assessing the difference between each dataset’s respective saturation results. This test is suitable15 because subjects are independent, and the measured differences are normally distributed (per the central limit theorem, sample sizes ≤ 30 are normally distributed). The formula used for the Unpaired T-Test is as follows, Figure 6:

**Figure 6.**
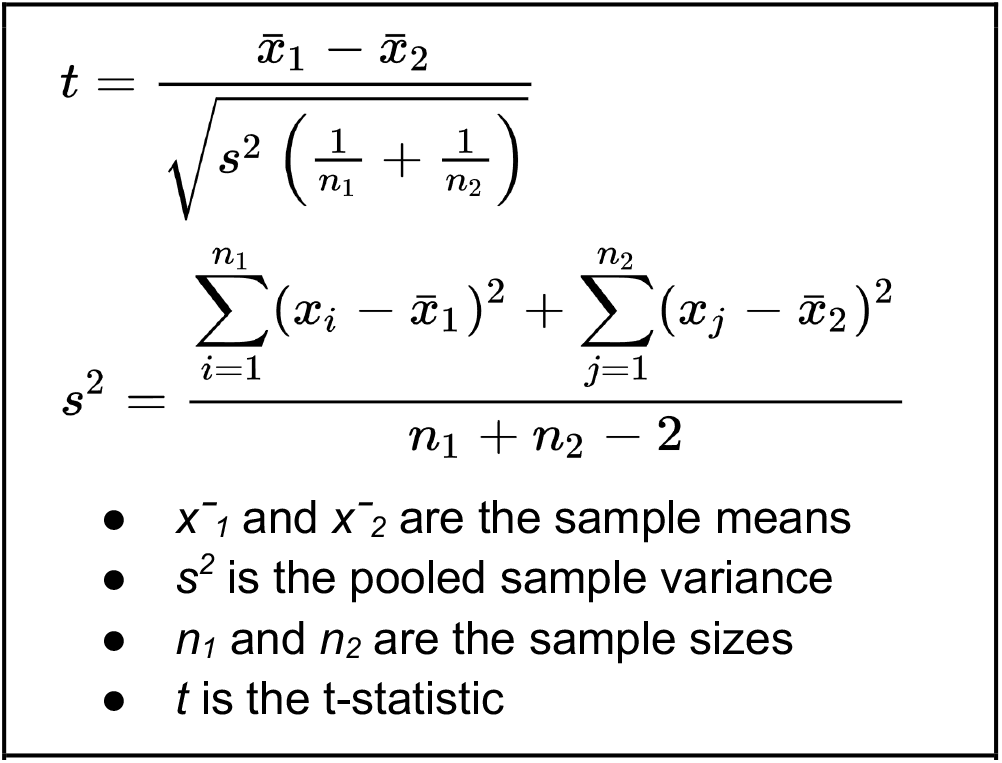
The formula used for the Unpaired T-Test.

Once uploaded, THRESHOLD outlines the result of your analyses, describing whether the differences between the saturation types in each data set yielded statistically significant differences. Additional relevant statistical measures, including the calculated p-value, are also listed. A more detailed description of the statistical analyses can also be exported with a (.txt) file of relevant calculated statistical measures for each of the saturation types.

### Furthered Analysis

Saturation data including saturation curves and the most saturated gene ascertained from the THRESHOLD tool were leveraged to facilitate further analyses and demonstrate broader use cases of THRESHOLD. g:Profiler (16) was utilized to conduct g:GOST functional profiling to understand the functional role of the most saturated genes identified. Proteinarium (17) was employed to construct protein-protein interaction networks, while Gephi (18) was utilized for visualizing these networks and applying various network parameters to better understand the critical pathways and communities driving cancer pathology. Moreover, network metrics such as modularity, separation testing (19), and Jaccard Similarity (20) were leveraged to understand patterns within and between the created protein-protein interaction networks.

### GUI Overview

THRESHOLD is present as a local GUI, Figure 7. When run, a home page facilitates a new session or statistical comparison of existing analyses. To investigate a new data set, a .txt file can be uploaded and relevant saturation parameters to export a graph of saturation supporting results. To perform a statistical comparison, upload two reference .txt files of THRESHOLD’s exported saturation data to assess whether there are statistically significant differences between samples to provide additional insights in user analyses.

**Figure 7.**
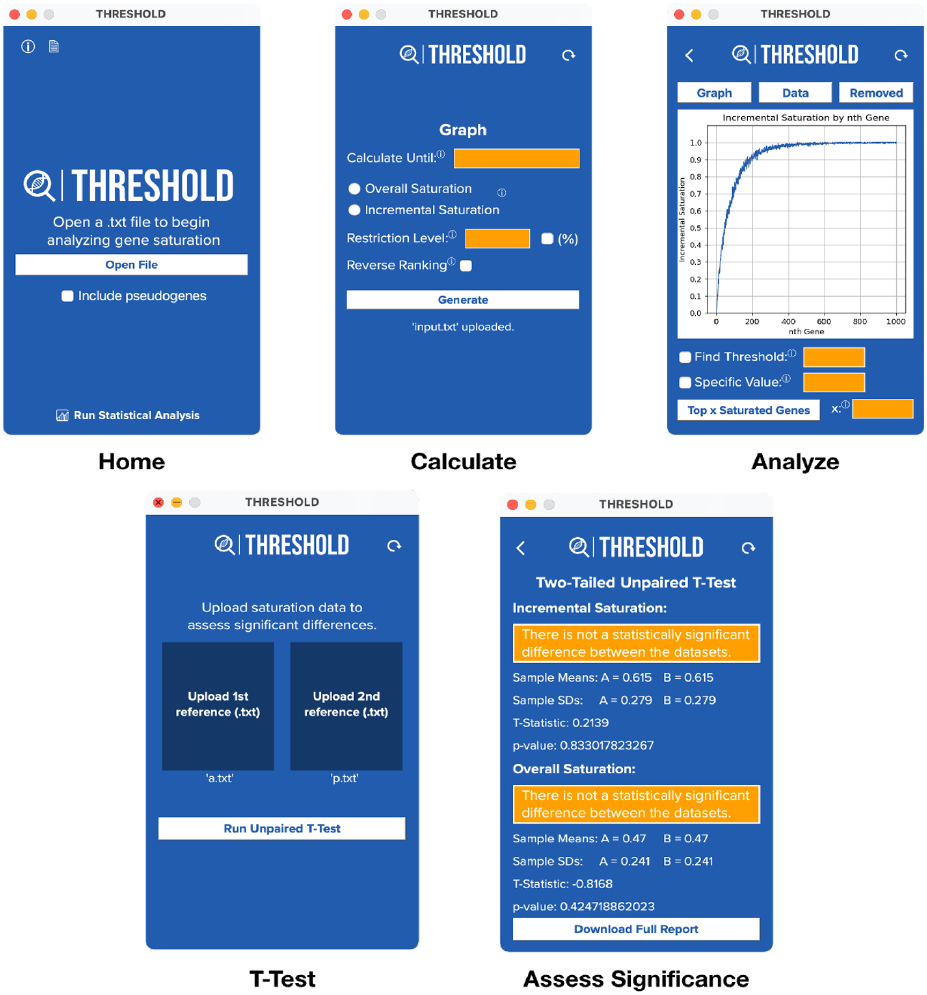
GUI Overview. Results.

### Validation Testing

Validation testing was conducted on a small test data set to verify the calculations outputted by THRESHOLD, Table 1. From our testing, the tool has demonstrated consistent accuracy, Table 2.

**Table 1.**
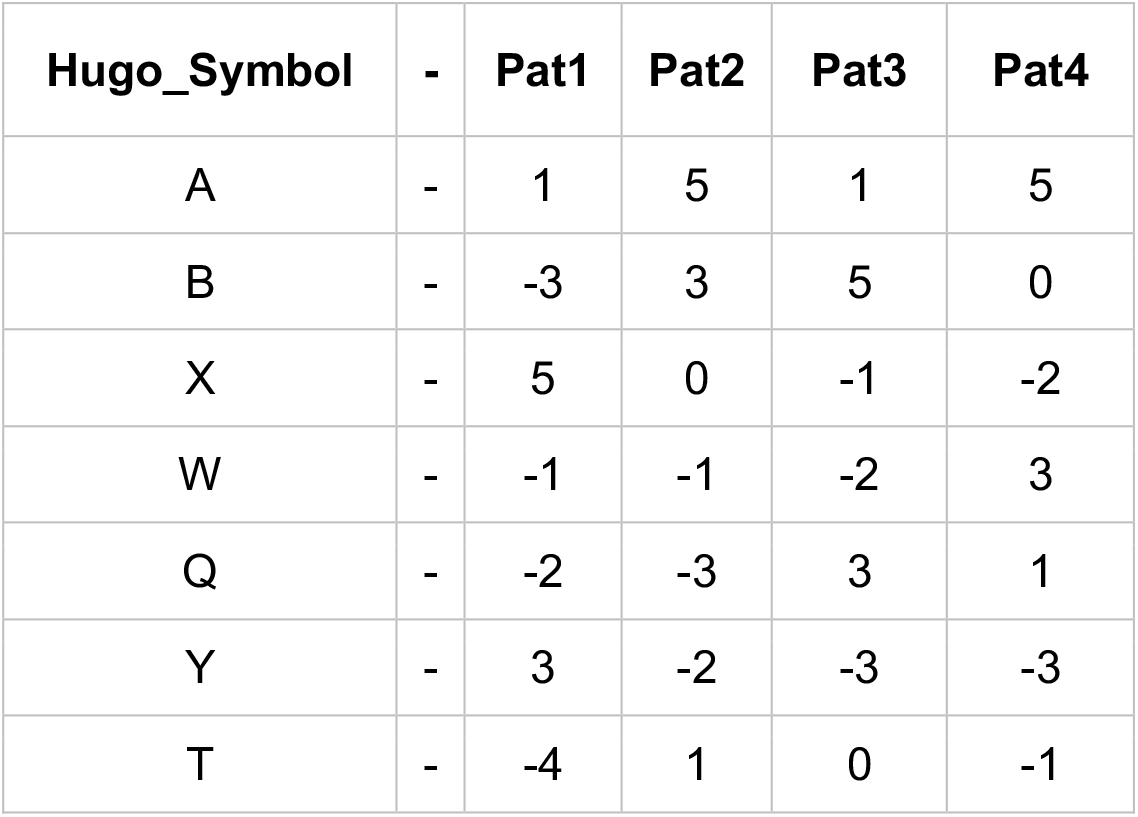
Saturation Test Dataset. An abbreviated data set of random “genes” represented by letters and corresponding z-scores representing expression was generated. The data set size was chosen to best permit and illustrate manual calculation of saturation per expanding computational complexity with increased dimensions. This data can then be used to calculate manual saturation values to verify THRESHOLD’s outputs.

**Table 2.** T-Test Test Dataset. Two arbitrarily generated saturation data references were generated to verify THRESHOLD’s statistical analysis outputs. The data sets mirror saturation data automatically exported by the THRESHOLD tool. Reference 1 and Reference 2 were then compared to corroborate calculated statistically significant differences and corresponding statistical parameters.

| *Saturation Data Reference 1* |  |  | *Saturation Data Reference 2* |  |  |
| --- | --- | --- | --- | --- | --- |
| **Nth Gene Included** | **Incremental Saturation** | **Overall Saturation** | **Nth Gene Included** | **Incremental Saturation** | **Overall Saturation** |
| 1 | 0.2 | 0.2 | 1 | 0.1 | 0.1 |
| 2 | 0.3 | 0.2 | 2 | 0.4 | 0.3 |
| 3 | 0.4 | 0.3 | 3 | 0.5 | 0.4 |
| 4 | 0.6 | 0.4 | 4 | 0.5 | 0.4 |
| 5 | 0.5 | 0.4 | 5 | 0.4 | 0.4 |
| 6 | 0.6 | 0.4 | 6 | 0.7 | 0.4 |
| 7 | 0.7 | 0.4 | 7 | 0.8 | 0.5 |
| 8 | 0.8 | 0.8 | 8 | 0.9 | 0.5 |
| 9 | 0.9 | 0.8 | 9 | 0.95 | 0.5 |
| 10 | 0.9 | 0.8 | 10 | 0.9 | 0.5 |

Given the following test dataset:

THRESHOLD passed verified calculations all the following tests:

-*Regular rank (level = 1, n = 5) - Reverse rank (level = 1, n = 5)*

-*Regular rank (level = 2, n = 5) - Reverse rank (level = 2, n = 5)*

-*Regular rank (level = 3, n = 5) - Reverse rank (level = 3, n = 5)*

-*Regular rank (level = 1% n = 5) - Reverse rank (level = 1% n = 5)*

-*Regular rank (level = 60%, n = 5) - Reverse rank (level = 60%, n = 5)*

-*Regular rank (level = 70%, n = 5) - Reverse rank (level = 70%, n = 5)*

Furthermore, we assessed THRESHOLD’s Paired T-Test function. We compared the tool’s outputted statistical metrics vs. our validated calculations to verify its results.

We compared the following two test data sets:

There was no difference found in the calculated mean differences, standard deviations, pooled standard deviation, standard error, and t-statistic. This validates THRESHOLD’s output statistical measures.

### Application of THRESHOLD Tool

#### 1) Bladder Urothelial Carcinoma Use Case

The THRESHOLD tool was used to elucidate insights into variable gene expression between bladder urothelial carcinoma patients. Patients were divided by cancer stage (II, III, and IV) and mRNA-seq data was analyzed to assess differences in saturation through Bladder Urothelial Carcinoma progression, (**Figure 8a**). Upon compiling each of the upregulated saturation curves **(Figure 8b)**, a statistically significant difference (p < 0.00001) was found between each of the upregulated overall curves, with progressive stages yielding heightened saturation values. These results were reciprocated in the top 10 most saturated genes from each of the respective stages, with the most saturated genes being present in growing proportions of patients by progressive stages (**Figure 8c**). Particularly, greater expression of cystatins including CST2 and CST4 which were the first and second most saturated upregulated genes, respectively, in Stage IV bladder urothelial carcinoma patients was observed through progressive stages. In addition, including COL10A1 and COL11A which were the third and fourth most saturated upregulated genes, respectively, in Stage IV bladder urothelial carcinoma patients were also observed through progressive stages. This trend is further supported by the greater observed Jaccard Similarity between adjacent stages (**Figure 8e)**, suggesting the progressive expression of certain critical saturated genes throughout bladder urothelial carcinoma stage progression. Moreover, functional profiling (g:GOST) performed on the top 10 most saturated genes in Stage IV cancer yielded significant major biological pathways and molecular functions relevant to Bladder Cancer progression (**Figure 8d)**. Most notably, these genes were implicated in cysteine-type endopeptidase inhibitor activity (p = 1.311×10^-4^) and endopeptidase inhibitor activity (p = 1.105×10^-2^) in addition to collagen degradation (p = 1.318×10^-4^) activity. The THRESHOLD tool was used to elucidate insights into expression profiles of bladder urothelial carcinoma patients based on datasets divided by sex **(Figure 9a)**. THRESHOLD found no statistically significant difference (p > 0.05) in the saturation between the male and female overall saturation curve (**Figure 9b**). Upon further investigation into the most saturated genes in each dataset, similar top 10 most saturated genes were identified regardless of dataset, including cystatins and collagen alpha one genes (**Figure 9c**). However, greater cystatin expression was observed in the female dataset, with CST2 and CST4 were present in 67% and 66% of female patients, respectively, juxtaposed to the male dataset in which CST2 and CST4 were present in 58% and 61% of male patients.

**Figure 8.**
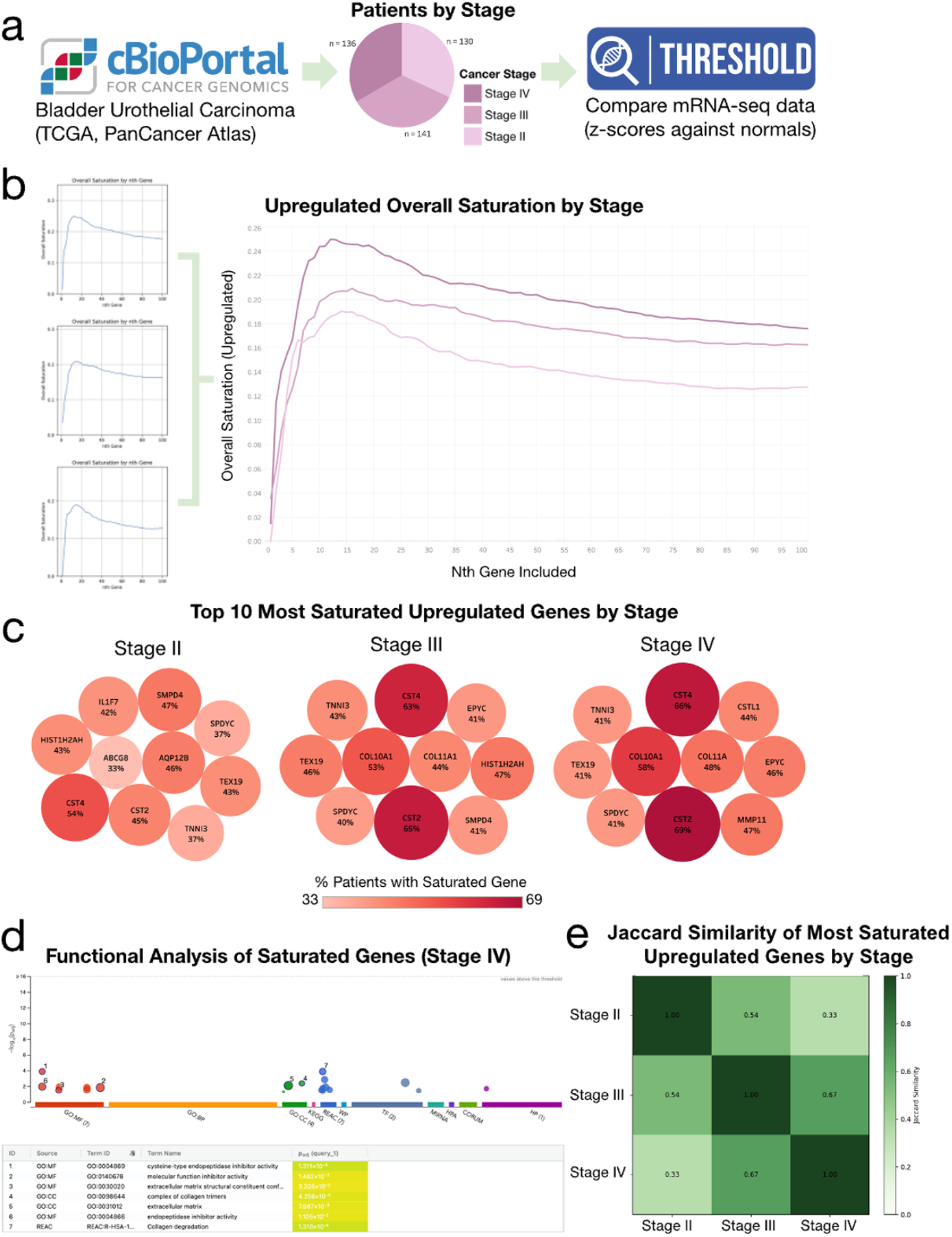
Bladder Urothelial Carcinoma Case Study. **a)** Analysis workflow. Datasets acquired from cBioPortal for Cancer Genomics, divided by cancer stage and mRNA-seq data were compared utilizing the THRESHOLD tool. **b)** Upregulated Overall Saturation of bladder urothelial carcinoma by Stage. Overall Saturation (restriction level 10%) was calculated for each of the cancer datasets by stage. Data was compiled into one graph elucidating differences in gene saturation by gene expression. **c)** Top 10 Most Saturated Upregulated Genes by Stage. The 10 Most Saturated Genes from each of the Bladder Urothelial Carcinoma datasets visualized indicating the percent of patients expressing the saturated gene within the nth gene rank included, 100 (**Figure 7b**). **d)** Functional Analysis of the Most Saturated Upregulated Genes for Stage IV bladder urothelial carcinoma. g:Profiler Functional profiling (g:GOST) performed on the top 10 most saturated gene (**Figure 7c**) from the Stage IV data set. **e)** Jaccard Similarity of Most Saturated Upregulated Genes by Stage. The top 10 saturated genes were compared using Jaccard Similarity to assess differences among high expression at each stage.

**Figure 9.**
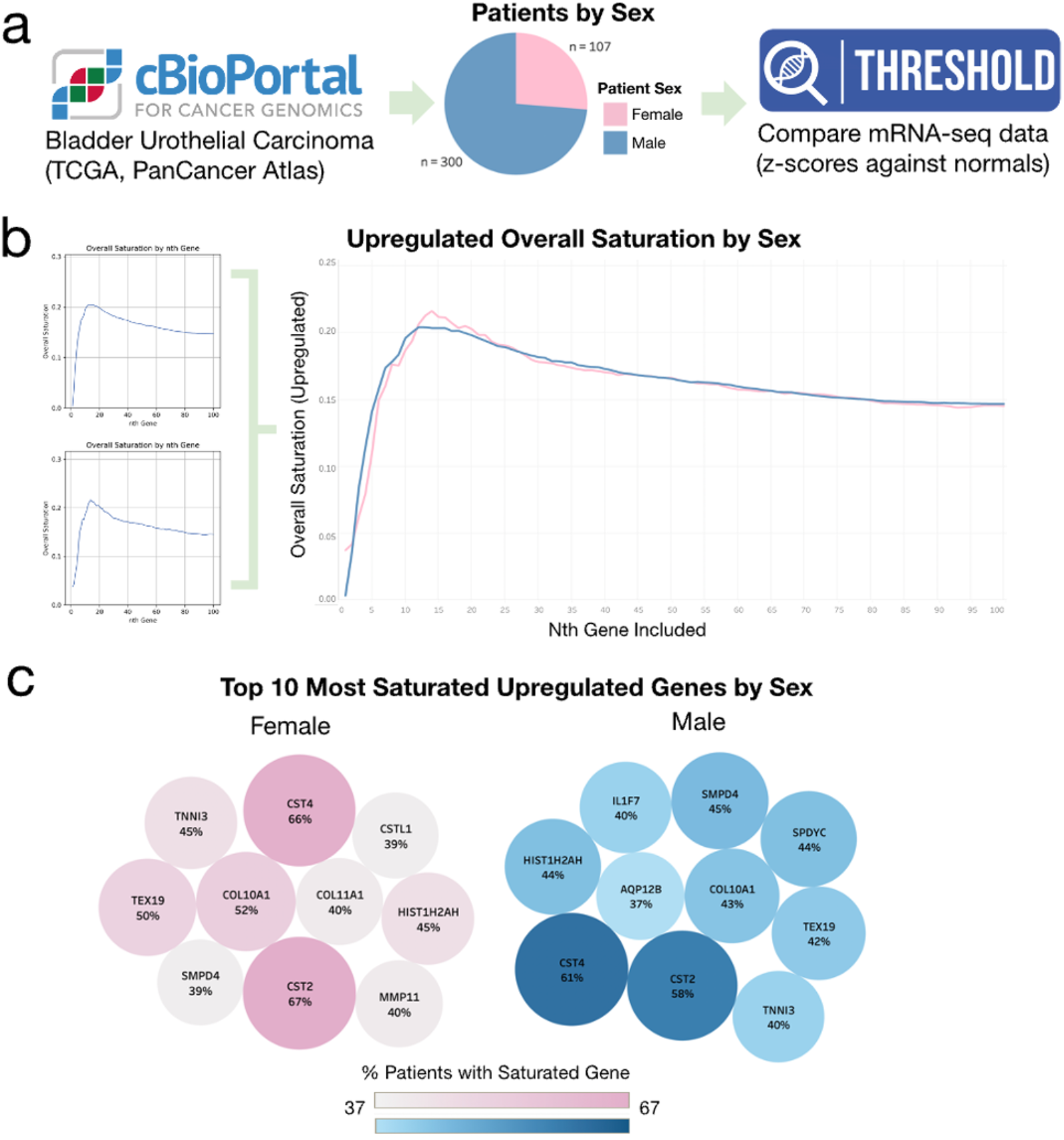
Bladder Urothelial Carcinoma Case Study Continued. **a)** Analysis workflow. Datasets acquired from cBioPortal for Cancer Genomics, divided by sex and mRNA-seq data were compared utilizing the THRESHOLD tool. **b)** Upregulated Overall Saturation of bladder urothelial carcinoma by sex. Overall Saturation (restriction level 10%) was calculated for each of the cancer datasets by stage. Data was compiled into one graph elucidating differences in gene saturation by gene expression. **c)** Top 10 Most Saturated Upregulated Genes by Stage. The 10 Most Saturated Genes from each of the Bladder Urothelial Carcinoma datasets visualized indicating the percent of patients expressing the saturated gene within the nth gene rank included, 100 (**Figure 6b**).

#### 2) Prostate Adenocarcinoma Use Case

To discern insights in the relationship between onset of prostate adenocarcinoma and gene expression, the THRESHOLD tool was leveraged to compare gene expression across patient samples clustered by diagnosis age. Patients were distributed into three age groups (41-57, 58-64, and 65-78) and compiled mRNA-seq data was analyzed to assess variations in saturation of different onsets of prostate adenocarcinoma **(Figure 10a)**. Comparative analysis of each of the overall upregulated saturation curves (**Figure 10b**) yielded a statistically significant difference (p < 0.0001), with younger onsets yielding heightened saturation values. The most saturated upregulated gene in all age-groups was SLC45A2, demonstrating progressive prevalence in younger age groups (**Figure 11a**). AMACR was also a top 10 most saturated gene in all age groups with progressive prevalence in younger age groups. However, all disease onsets demonstrated similar degrees of overall downregulated saturation, with high degrees of cohesion in all age groups (**Figure 10c**). The most saturated downregulated genes driving this high expression include olfactory receptor (OR) genes including OR9Q1 and OR2AT4 in addition to a histone regulating gene, HIST1H4F present in all age groups’ top 10 most saturated downregulated genes (**Figure 11b**). Heightened levels of Jaccard Similarity between top 10 most saturated downregulated genes in each age group (**Figure 11d**) further support the notion of a high degree of downregulated gene saturation in prostate adenocarcinoma. Functional profiling (g:GOST) performed on the top 100 most saturated upregulated and downregulated genes (**Figure 10d, Figure 10e**) both yielded significant olfactory receptor activity, particularly in the downregulated gene set (p = 2.074×10^-20^). Moreover, the upregulated gene set indicated relevance in histidine synthase activity (p = 1.061×10^-2^) in addition to G protein-coupled receptor activity (p = 1.997×10^-3^)

**Figure 10.**
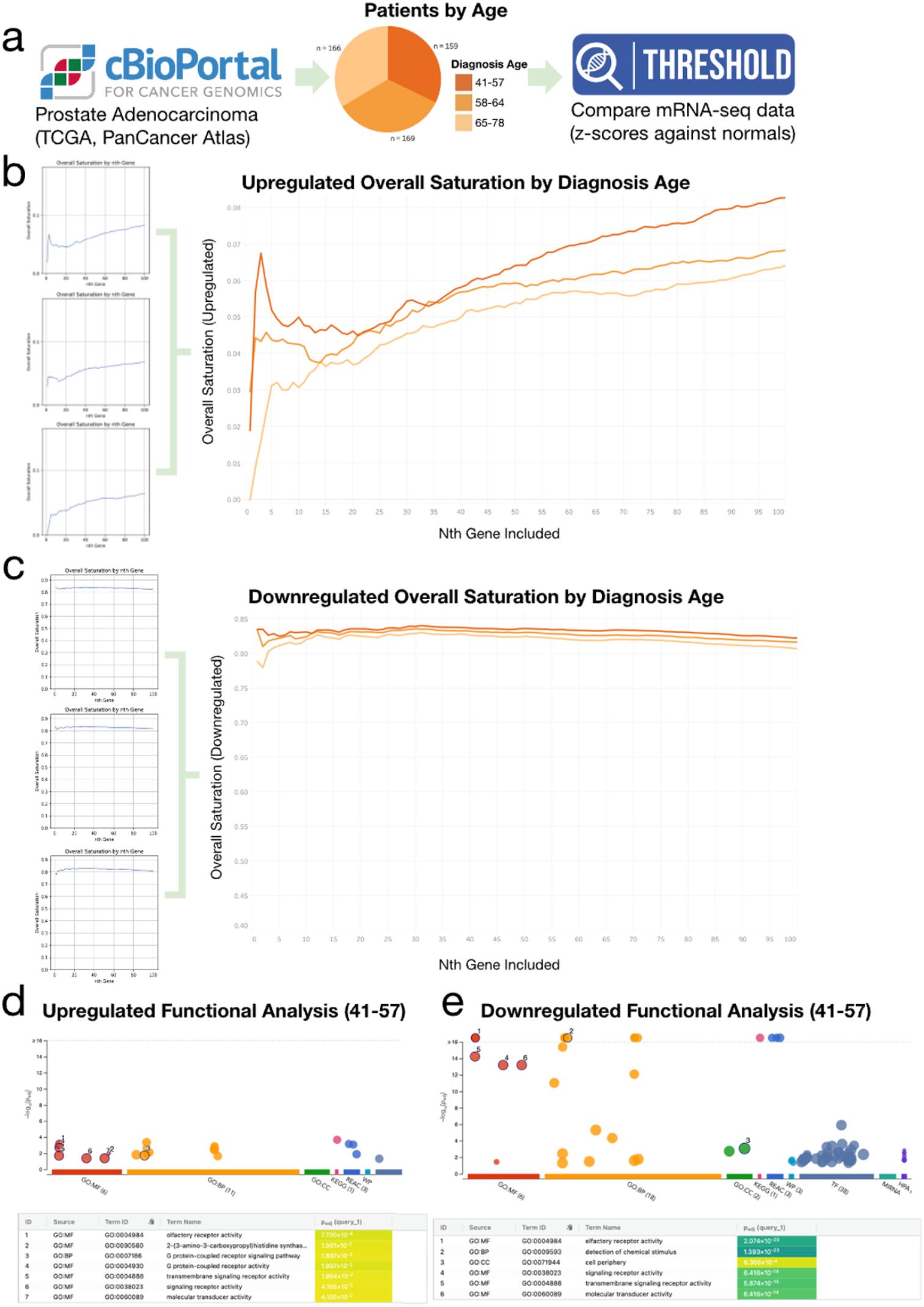
Prostate Adenocarcinoma Case Study. **a)** Analysis workflow. Datasets acquired from cBioPortal for Cancer Genomics, divided by diagnosis age and mRNA-seq data were compared utilizing the THRESHOLD tool. **b)** Upregulated Overall Saturation of Prostate Adenocarcinoma by Diagnosis Age. Overall Saturation (restriction level 10%) was calculated for each of the cancer datasets by age. Data was compiled into one graph elucidating differences in gene saturation by gene expression. **c** Downregulated Overall Saturation of Prostate Adenocarcinoma by Diagnosis Age. Overall Saturation (restriction level 10%) was calculated for each of the cancer datasets by age. Data was compiled into one graph elucidating differences in gene saturation by gene expression. **d)** Functional Analysis of the Most Saturated Upregulated Genes for Diagnoses between 41 and 57 in Prostate Adenocarcinoma. g:Profiler Functional profiling (g:GOST) performed on the top 10 most saturated upregulated genes (**Figure 7f**) from the diagnosis age 41-57 data set. **e)** Functional Analysis of the Most Saturated Downregulated Genes for Patients Diagnoses between the ages of 41 and 57 in Prostate Adenocarcinoma. g:Profiler Functional profiling (g:GOST) performed on the top 10 most saturated downregulated genes (**Figure 7h**) from the diagnosis age 41-57 data set.

**Figure 11.**
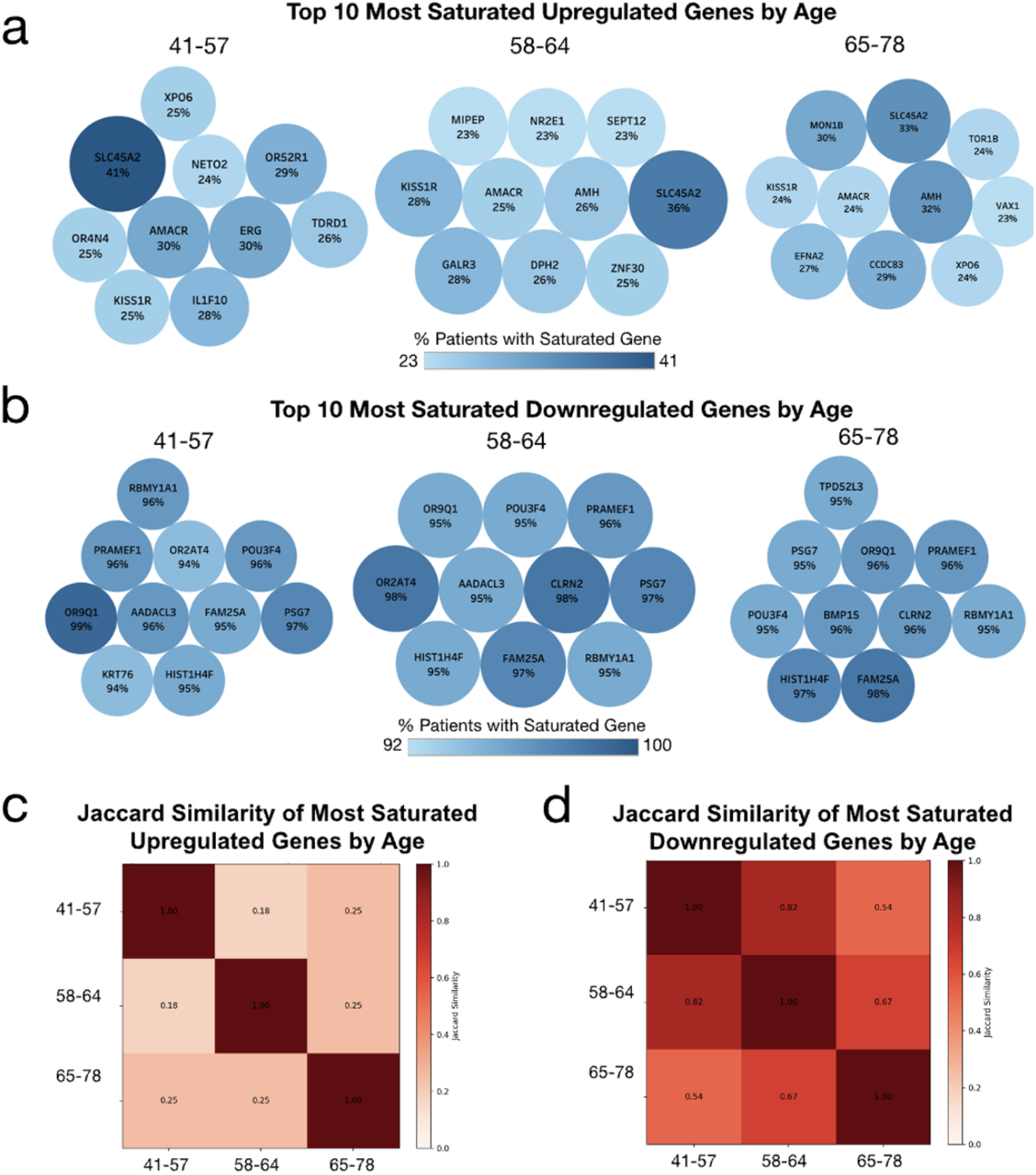
Prostate Adenocarcinoma Case Study Continued. **a)** Top 10 Most Saturated Upregulated Genes by Diagnosis Age. The 10 most saturated upregulated genes from each of the Prostate Adenocarcinoma datasets are visualized indicating the percent of patients expressing the saturated gene within the nth gene rank included, 100 (**Figure 7b**). **b)** Top 10 Most Saturated Downregulated Genes by Diagnosis Age. The 10 most saturated downregulated genes from each of the Prostate Adenocarcinoma datasets are visualized indicating the percent of patients expressing the saturated gene within the nth gene rank included, 100 (**Figure 7c**). **c)** Jaccard Similarity of Most Saturated Upregulated Genes by Diagnosis Age. The top 10 saturated upregulated genes were compared using Jaccard Similarity to assess differences among high gene expression at each stage. **d)** Jaccard Similarity of Most Saturated Downregulated Genes by Diagnosis Age. The top 10 saturated downregulated genes were compared using Jaccard Similarity to assess differences among low gene expression at each stage.

## Discussion

The THRESHOLD’s results demonstrate the numerous valuable analyses and insights into disease as a novel genomics tool. Through its results, we have validated the tool’s capacity for patient stratification and biomarker and drug targeting identification and found new areas for researchers to explore further. Our bladder urothelial carcinoma analyses (**Figure 8**) indicate that the THRESHOLD tool demonstrated a strong capacity to analyze gene expression underlying disease progression. THRESHOLD highlighted heightened unregulated gene saturation in progressive stages, suggesting greater gene cohesion as cancer develops within patients (**Figure 8b**). Further insights permitted the identification of the most critical genes driving this growth in saturation, including the identification of collagen alpha 1 genes and cystatins (**Figure 8c**). Collagen alpha 1 genes including COL10A1 identified as the third most saturated gene in Stage IV bladder. Functional analysis of COL10A1 and co-expressed genes has indicated significant relevance in ECM-receptor interaction, protein digestion and absorption, and PI3K-AKT signaling pathways. The COL10A1 gene has also been implicated as a valuable prognostic and predictive biomarker (21) in bladder cancer with increased COL10A1 expression being related to poor overall patient survival.

Cystatins including CDC2 and CDC4 were also implicated as leading drivers of gene saturation progression in bladder urothelial carcinoma, representing the first and second most saturated genes in Stage IV Bladder Cancer B (**Figure 8c**). Prior literature validates these results, demonstrating that within patients with elevated cystatin expression (22) in urine such as Cystatin B, there was a short mean time to disease recurrence and in grade/stage progression of Transitional Cell Carcinoma (Urothelial Carcinoma). These results again demonstrate THRESHOLD’s utility in identifying novel predictive biomarkers implicated in disease progression, grade, and recurrence.

Moreover, the THRESHOLD tool demonstrated insights in evaluating differences in gene expression by clinical features such as age of diagnosis in the prostate adenocarcinoma datasets (**Figure 10**). THRESHOLD demonstrated heightened upregulated gene saturation in samples with earlier diagnosis ages, suggesting greater gene cohesion in patients with earlier onsets of prostate adenocarcinoma (**Figure 10b**). The most saturated genes driving this trend included SLC45A2 and AMACR, which demonstrated heightened saturation in younger ages (**Figure 11a**). This pair is significant as scientific literature has implicated the two genes as major fusion transcripts underlying prostate adenocarcinoma. In a study (23) of eight fusion transcripts including SLC45A2-AMACR, it was demonstrated that 91% of patients positive for any of the fusion transcripts experienced recurrence, metastasis, or prostate adenocarcinoma-associated death even after radical prostatectomy as compared to only 37% of patients not carrying the fusion transcripts. This suggests THRESHOLD could be used as a tool to identify fusion transcripts implicated in disease severity.

Additionally, Prostate Adenocarcinoma expression data yielded significantly downregulated gene saturation in all diagnosis age groups, suggesting pronounced suppression of several genes (**Figure 10c**). The most downregulated saturated genes demonstrated an intriguing pattern of olfactory receptor genes such as OR9Q1 and OR2AT4 (**Figure 11b**), and functional analyses also revealed significant activities concerning olfactory receptors (**Figure 10e**). In prostate adenocarcinoma specifically, the olfactory receptor gene OR51E2 has been implicated (24) in activating ERK1/2 via the Gβγ-PI3Kγ-ARF1 pathway elucidating important insights into characteristic MAPK hyper-activation. These results validate THRESHOLD’s relevance in elucidating critical gene pathways underlying differential progression or manifestation of disease.

In the supplementary, colon adenocarcinoma analyses, we demonstrated THRESHOLD’s utility in elucidating novel drug targets among developed networks of highly saturated genes (**Figure S1**). Upon generation of an overall upregulated saturation curve (**Figure S1b**), the most saturated genes (**Figure S1c**) were extracted for network analyses and functional profiling. Of these network community hub genes and most saturated genes included actins, such as ACTB and ACTG1. ACTB and ACTG1 have been implicated as a significant biomarkers and regulator of tumorigenicity in numerous cancers including hepatomas, renal cell carcinoma, and colon adenocarcinoma as we investigated. ACTB regulates (25) F-actins which play roles in chemoresistance and increased cell proliferation. Moreover, ACTB can increase membrane protrusions, focal adhesions, and myosin activity, which increase tumor migration. In parallel, ACTG1 increases (25) cell proliferation through the mitochondrial apoptotic pathway, Warburg effect, upregulation of CDKs, and ROCK signaling pathways. Collectively, these roles suggest actins including ACTB and ACTG1 could serve as biomarker and drug targeting candidates in several cancers including colon adenocarcinoma.

Extended networks were generated from these most saturated genes and modularity testing yielded compelling communities of interest (**Figure S1d, Figure S1f**). This included a community around HSPA8, the most interconnected node with degree ten (**Figure S1g**). HSPA8 (also known as Hsc70) (26-30) has been implicated as a key driver of cell proliferation (31) under many conditions, with its absence inhibiting the growth of tumors and via apoptosis and cell cycle arrest. As indicated by the network, HSPA8 also interacts with MAPK1 via its broader community (**Figure S1f**), a signaling pathway under intense investigation for its relevance in a multitude of cancers. For these reasons, HSPA8 has been recognized as a biomarker candidate for the early detection and diagnosis of cancers including endometrial carcinoma (32) and colon adenocarcinoma (33). This gene, among several other network hub genes including YWHAZ and EEF1A1 could serve as novel drug and therapeutic targets, representing critical hubs underlying colon adenocarcinoma pathology.

THRESHOLD evaluated if there were differences in expression among breast invasive carcinoma patients undergoing or not undergoing radiation treatment (**Figure S2**). After separation of patient populations and comparison of individually calculated incremental saturation curves, there was no significant difference between either curve (**Figure S2b, Figure S2c**). This suggests that the radiation treatments administered did not significantly impact gene expression between the sample populations and was further supported by networks developed via the most upregulated genes that were highly overlapping in the interactome with an sAB of -0.84 (**Figure S3f**). Additionally, similarly to prostate adenocarcinoma, olfactory receptor genes were highly downregulated as derived from the downregulated incremental saturation curve (**Figure S2e**). Functional analysis also implicated olfactory activity as a highly significant (p = 2.310×10^-30^) function of the top 100 most saturated downregulated genes. Several of these olfactory receptor genes have been implicated in cancer metastasis, invasion, and proliferation, through signaling pathways such as NF-κB/STAT (34) in breast cancers. Furthermore, modularity testing of the protein-protein interaction networks formed from the top 100 most upregulated saturated in each treatment group (**Figure S3b, Figure S3d**) revealed significant insights into biomarker candidates and drug targets for new therapies. For example, overexpression of highly interconnected hub genes such as RBBP7 regulating chromatin metabolism has been associated with poor overall patient survival (35) in cancers such as esophageal squamous cell carcinoma, being implicated with enhanced tumor migration and invasion. Additionally, in adjacent network communities, MAPKAPK2 (MK2) was present and has significant relevance as a breast cancer biomarker and therapeutic target. The p38MAPK-MK2 signaling pathway contributes to the stimulation of triple-negative breast cancer tumorigenesis by promoting AP1 activity, (36) associated with aggressive cancer manifestations. In all, the breast invasive carcinoma use case demonstrated utility in identifying drug targets and biomarkers, in addition to establishing where homogeneity may exist between samples of varying clinical features.

## Conclusion

The THRESHOLD bioinformatics tool serves as a robust and innovative approach for deciphering shared gene expression patterns across patient populations. Its dual functionalities, incremental saturation and overalls saturation, provide both granular and overarching perspectives on gene expression and facilitate relevant comparative analyses. This makes THRESHOLD not only pivotal for disease research, drug repurposing studies, and personalized medicine but also for understanding the stepwise changes in gene expression as diseases progress or respond to treatment. Moreover, adapting the THRESHOLD to focus on suppressed genes could further extend its utility in exploring downregulated or silenced pathways in various conditions. Collectively, the tool offers significant potential in advancing our understanding of molecular signatures, streamlining patient stratification, and fostering the development of tailored therapeutic interventions.

The THRESHOLD tool is freely available at: https://github.com/alperuzun/THRESHOLD.

