## Supplemental File 1 for "THRESHOLD: A Comprehensive Transcriptomic Analysis Tool for Evaluating Gene Saturation and Impact in Disease Progression"

### Supplementary Results

#### 1) Colon Adenocarcinoma Case Study

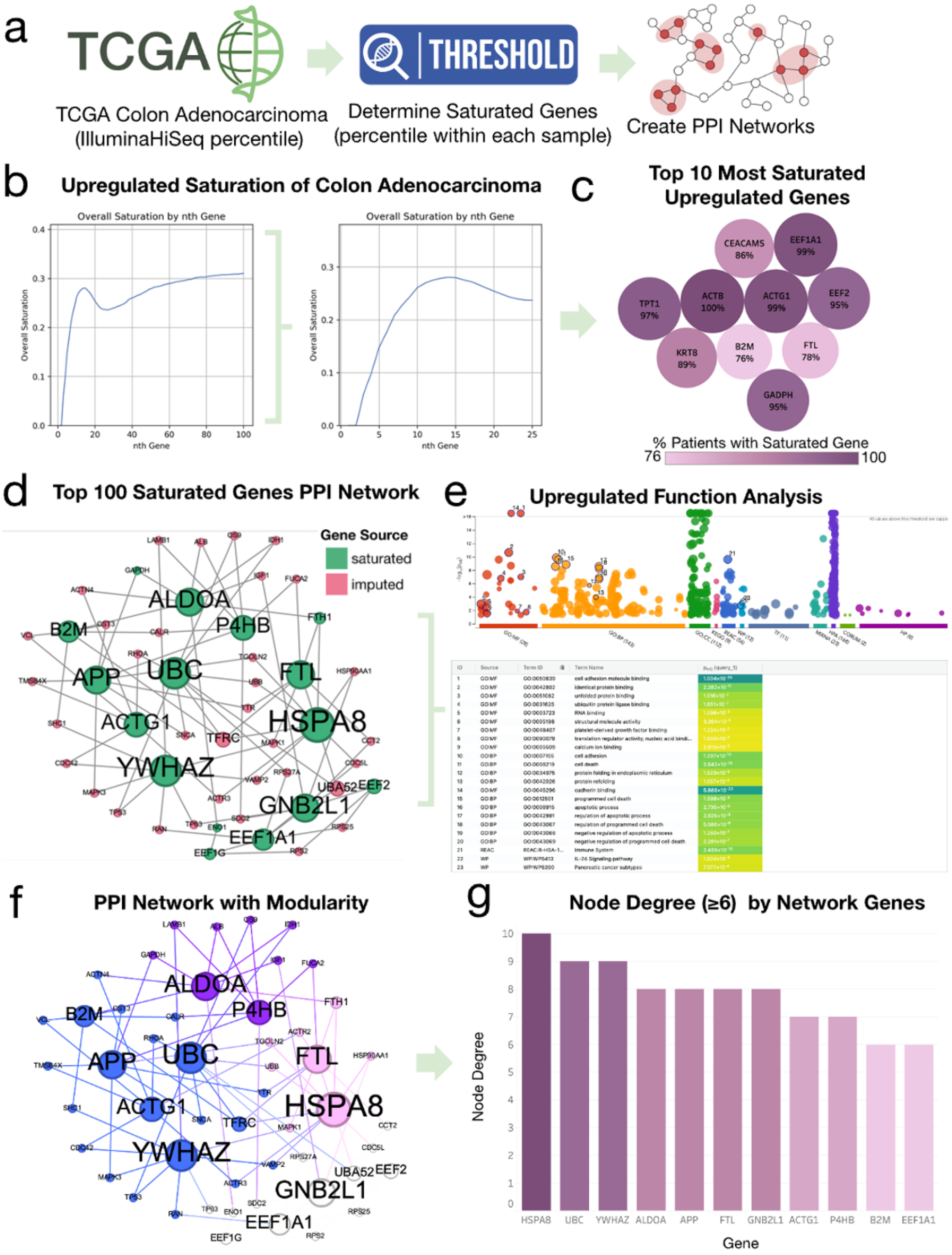


**Figure S1. Colon Adenocarcinoma Case Study. a** Analysis workflow. Datasets acquired from TCGA database via UCSC Xena, the THRESHOLD tool was used to compare mRNA-seq data to create PPI networks to elucidate potential drug targets. **b** Upregulated Overall Saturation of Colon Adenocarcinoma. The first curve illustrates overall saturation calculated (restriction level 40%) up to 100 genes while the second curve focuses on a pronounced maximum up to 25 genes. **c** Top 10 Most Saturated Upregulated Genes. The top 10 most saturated upregulated genes from the colon adenocarcinoma dataset visualized indicating the percent of patients expressing the saturated gene within the nth gene rank included (**Figure S1b**). **d** Top 100 Saturated Upregulated Gene Protein-Protein Interaction Network via Proteinarium. The top 100 saturated upregulated genes (**Figure S1b**) were used to create PPI networks including genes imputed into relevant pathways via Proteinarium. **e** Functional Analysis of Most Saturated Genes. g:Profiler Functional profiling (g:GOST) performed on the top 100 most saturated upregulated genes to elucidate potential pathways underlying disease pathology. **f** Refined Protein-Protein Interaction Networks with Modularity. Protein interaction data was leveraged to calculate and visualize the modularity of the network and communities within the overall PPI network (**Figure S1d**). **g** Node Degree ≥6 by Network Genes. Proteins of degree 6 (6 or more network interactions) were extracted from the network data to elucidate most interconnected hub genes as potential drug targets.

We gathered within-sample percentile data for colon adenocarcinoma to explore THRESHOLD's capacity to facilitate the creation of protein-protein interaction networks to elucidate the most essential biological pathways underlying cancer pathology and to garner insights for relevant drug targets. Upregulated overall saturation for the data set was calculated and yielded a pronounced crest within the first 25 ranked genes, suggesting cohesion of the most upregulated genes across samples (**Figure S1b**). The 10 most saturated upregulated genes from this analysis included actins such as ACTB and ACTG1, in addition to elongation factors such as EEF1A1 and EEF2 (**Figure S1c**). The extended list of the 100 most saturated upregulated colon adenocarcinoma genes was used to create a protein-protein interaction network, with relevant genes imputed to complete the relevant network interactions (**Figure S1d**). Furthered modularity testing yielded analysis within the networks, elucidating hub genes and communities underlying specific pathways (**Figure S1f**). The most interconnected hub genes included HSPA8, UBC, and YWHAZ (**Figure S1g**), each implicated in significant connections within each of their respective communities. Functional profiling of these rigorous networks yielded significant insights in the underlying biological pathways, including numerous implications in the regulation of apoptosis including negative regulation of apoptotic process ($p = 1.280*10^{-7}$) and regulation of apoptotic process ($p = 2.626*10^{-9}$) in addition to binding in cell adhesion molecule binding ($p = 1.004*10^{-25}$) (**Figure S1e**).

### 2) Breast Invasive Carcinoma Case Study

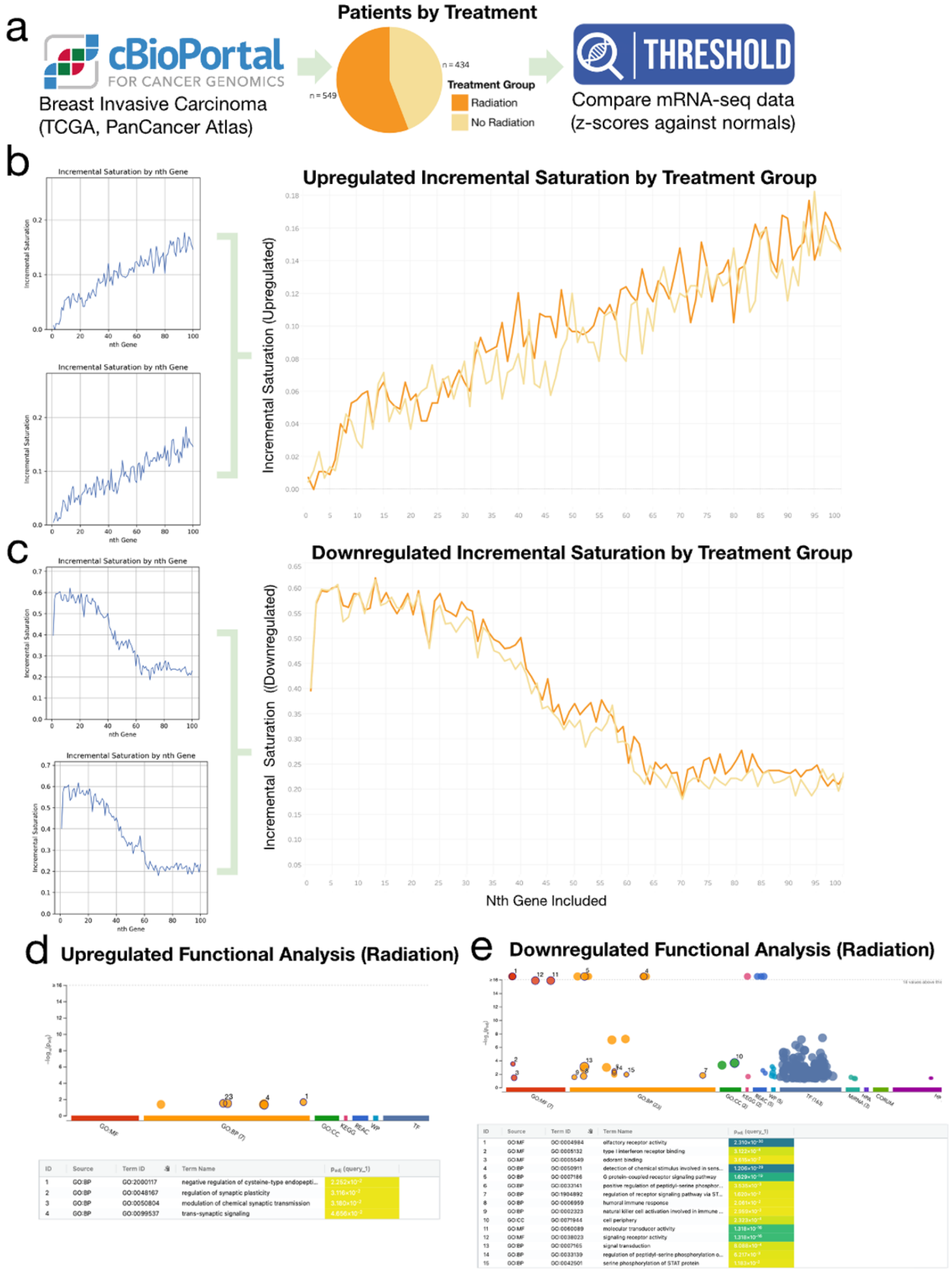


**Figure S2. Breast Invasive Carcinoma Case Study. a** Analysis workflow. Datasets acquired from cBioPortal for Cancer Genomics, divided by treatment group and mRNA-seq data were compared utilizing the THRESHOLD tool. **b** Upregulated Incremental Saturation of Breast Invasive Carcinoma by Diagnosis Age. Incremental Saturation (restriction level 10%) was calculated for each of the cancer datasets by treatment group. **c** Downregulated Incremental Saturation of Breast Invasive Carcinoma by Treatment Group. Incremental Saturation (restriction level 10%) was calculated for each of the cancer datasets by treatment group. Data was compiled into one graph elucidating differences in gene saturation by gene expression. **d** Functional Analysis of Most Saturated Upregulated Genes (Radiation). g:Profiler Functional profiling (g:GOST) performed on the top 100 most saturated upregulated genes (**Figure S2b**) from the radiation treatment group. **e** Functional Analysis of Most Saturated Downregulated Genes (Radiation) g:Profiler Functional profiling (g:GOST) performed on the top 100 most saturated downregulated genes (**Figure S2c**) from the radiation treatment group.

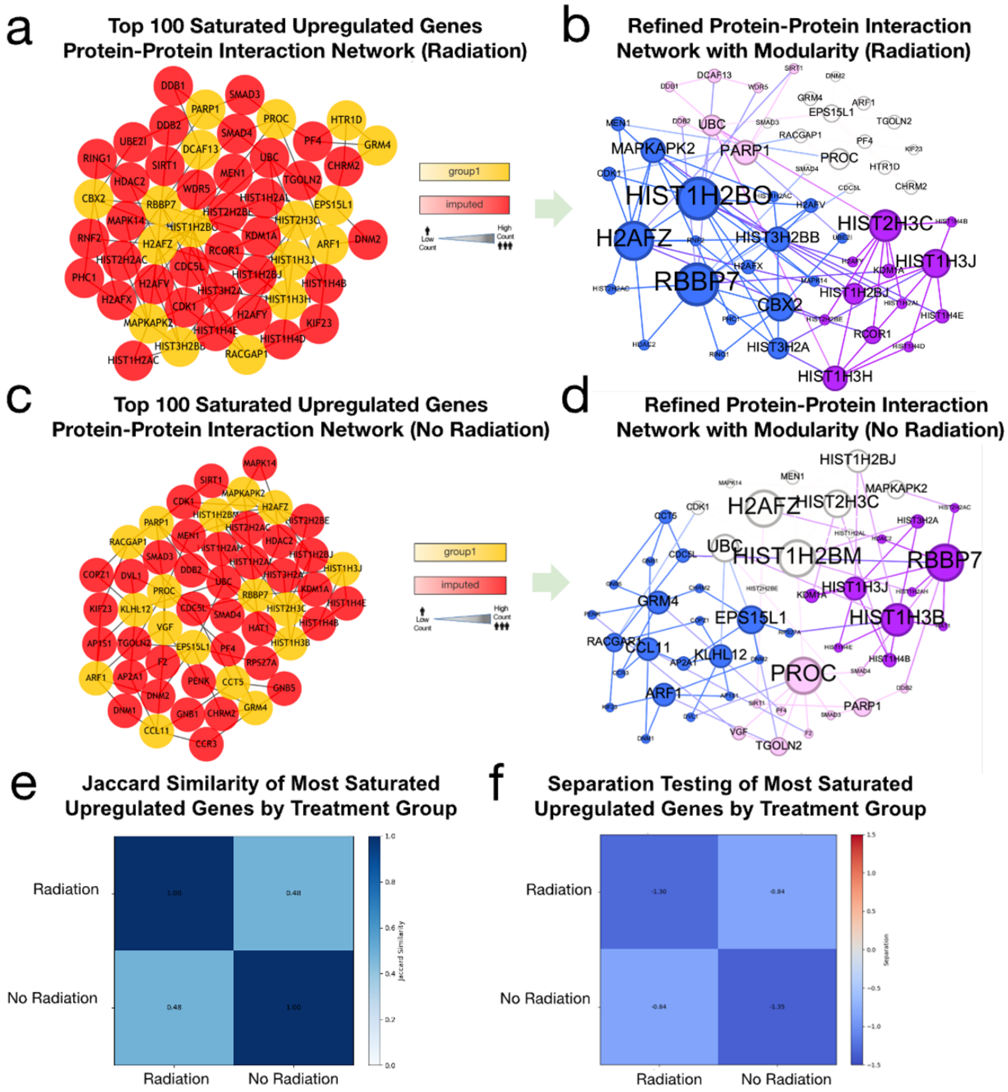


**Figure S3. Breast Invasive Carcinoma Case Study Continued. a** Top 100 Saturated Upregulated Gene Protein-Protein Interaction Network via Proteinarium (Radiation treatment group). The top 100 saturated upregulated genes from the radiation treatment group (**Figure S2b**) were used to create PPI networks including genes imputed into relevant pathways via Proteinarium. **b** Refined Protein-Protein Interaction Networks with Modularity (Radiation treatment group). Protein interaction data was leveraged to calculate and visualize the modularity of the network and communities within the overall PPI network (**Figure S3b**). **c** Top 100 Saturated Upregulated Gene Protein-Protein Interaction Network via Proteinarium (No radiation treatment group). The top 100 saturated upregulated genes from the no radiation treatment group (**Figure S2c**) were used to create PPI networks including genes imputed into relevant pathways via Proteinarium. **d** Refined Protein-Protein Interaction Networks with Modularity (No radiation treatment group). Protein interaction data was leveraged to calculate and visualize the modularity of the network and communities within the overall PPI network (**Figure S3d**). **e** Jaccard Similarity of Most Saturated Upregulated Genes by Treatment Group. The saturated genes (and relevant imputed genes) were compared using Jaccard Similarity to assess differences between high gene expression in each treatment group. **f** Separation Testing of Most Saturated Upregulated Genes by Treatment Group. The protein-protein interaction networks underwent separation testing to elucidate similarities or differences between pathway functions via distance between two networks (**Figure S3e**, **Figure S3f)** in the interactome.

The THRESHOLD tool was used to elucidate insights into variable gene expression between breast invasive carcinoma patients. Patients were divided by treatment group (radiation treatment and no radiation treatment) and mRNA-seq data was analyzed to assess differences in saturation between treatment groups. The upregulated incremental saturation curves were calculated and compiled into one graph. There was not a statistically significant difference between the treatment groups ($p > 0.05$), demonstrating similar gene saturation between both curves (**Figure S2b)**. Functional profiling (g:GOST) performed on the top 100 most saturated genes in the radiation treatment group yielded significant major biological pathways and molecular functions relevant to breast invasive carcinoma. These saturated genes were implicated in the negative regulation of cysteine-type endopeptidase activity among other cellular signaling roles (**Figure S2d**). Similarly, the downregulated incremental saturation curves did not yield statistically significant results between the treatment groups ($p > 0.05$), again demonstrating similar gene saturation between both curves (**Figure S2c**). Both downregulated incremental saturation curves demonstrated relatively heightened incremental saturation among the initial ranked genes, suggesting high gene saturation among the most downregulated genes. The downregulated genes thus yielded more significant g:Profiler Functional profiling (g:GOST) results, including implications in olfactory receptor activity ($2.310*10^{-30}$), G protein-coupled report signaling pathways ($1.629*10^{-19}$), among other signaling and signal transduction pathways (**Figure S2e**). The top 100 upregulated saturated from both the radiation and no radiation treatment groups were then used to create protein-protein interaction networks, including imputed genes in relevant pathways (**Figure S3a**, **Figure S3c**) to elucidate insights to more detailed pathways underlying breast invasive carcinoma pathology. These networks were then evaluated for modularity, distinguishing between communities within the broader networks. In the radiation treatment group protein-protein interaction networks, two separate communities were found around histidine hub genes including HISTH2BO and HIST2HC3 (**Figure S3b**). Other large hub genes were identified including MAPKAPK2, PROC, and PARP1. In the no radiation treatment group network, two separate communities were also found around histidine hub genes including HIST1H2BM and HST1H3B (**Figure S3d**). Other large hub genes were also identified including EPS15L1, PROC, and UBC. While there were some differences in the specific genes within the two networks as indicated by Jaccard Similarity (**Figure S3e**), the overall function of the networks is similar as suggested by their relatively negative sAB value -0.84 indicating they overlap in the interactome (**Figure S3f**).
